# First direct quantification of floral handling costs in bees

**DOI:** 10.1101/2025.09.15.675838

**Authors:** Natacha Rossi, Mario Vallejo-Marín, Elizabeth Nicholls

## Abstract

Floral handling can be energetically costly for bees, yet these costs are rarely measured. We provide the first direct quantification of the metabolic cost of floral buzzing in bumblebees and evaluate its ecological significance. Using flow-through respirometry synchronised with laser vibrometry, we measured carbon dioxide production during buzzing by *Bombus terrestris* and compared it with flight take-off, which is powered by the same thoracic muscles. Buzzing required high muscular effort, with ∼0.10 J per event and mass-specific power ∼293 W kg⁻¹, and overall costs comparable to take-off because buzzing bouts are longer even though the metabolic rate is lower. Absolute metabolic rate increased with body mass, whereas intertegular span did not, implying that transient load rather than structural size better explains energetic demand in short, high-intensity behaviours. Metabolic traits were repeatable within individuals, and colony identity explained additional variance, consistent with genetic or shared environmental effects. Converting costs to nectar equivalents showed that buzzing required slightly more nectar than take-off and that requirements rose as nectar sugar concentration declined. We conclude that floral buzzing is a major, previously unquantified component of bee energy budgets that is likely to shape nectar supplementation, flower sequencing, and plant–pollinator interaction strength.

## Introduction

Most work on bee foraging emphasises time/cognitive and travel costs (1–8), yet floral handling, particularly pollen collection, also requires muscular work that can reduce visit profitability (9–12). Because pollen is a non-energetic reward for adult bees, typically collected to provision brood rather than fuel activity, and often requires mechanically complex extraction, these costs may not be offset by immediate energetic gain (13). Therefore, handling costs could have important consequences for bee foraging efficiency, plant fitness, and plant–pollinator interaction networks (14,15).

A particularly striking example is floral sonication, or buzz pollination, where bees vibrate flowers to extract pollen from poricidal anthers, a floral adaptation that restricts pollen access by enclosing grains within tube-like structures (15–17). This strategy is widespread, found in an estimated 20,000 flowering plant species across diverse taxa (e.g. Solanaceae, Melastomataceae) (14,18). To access these rewards, bees such as bumblebees use their indirect flight muscles (IFMs) to generate high-frequency thoracic vibrations that forcibly eject pollen through small apical pores (14,15,19). This behaviour has been recorded in more than 400 species, across 74 of the 508 recognised bee genera (20), indicating that vibratile pollen collection is both phylogenetically widespread and ecologically significant (14,21). Buzzing is often performed on flowers that offer no nectar reward and buzz frequencies can exceed those of flight, suggesting high muscular demands (14,22). Moreover, buzzing behaviour is modulable: bees can adjust buzz duration based on reward availability or experience (21,23,24), indicating a potentially plastic response to energetic or mechanical constraints.

Despite its prevalence and presumed cost, the energetics of floral buzzing have never been directly quantified. While the metabolic demands of flight are well-characterised and rank among the most energetically expensive behaviours on a mass-specific basis (25–27), floral buzzing remains unmeasured. Because both behaviours are powered by the same indirect flight muscles, flight provides a useful reference point for evaluating buzzing cost. Although buzzing may involve higher frequency and acceleration than flight (22), it lacks the aerodynamic and inertial constraints of lift generation. Whether this makes buzzing more or less metabolically costly than flight is unknown.

Here, we present a novel, integrative approach to quantifying the metabolic cost of floral buzzing, combining flow-through respirometry, laser vibrometry, and video recording to synchronise real-time CO_2_ production with distinct behavioural events. We apply this method to the buff-tailed bumblebee (*Bombus terrestris*) and compare buzzing to flight take-off, a high-intensity behaviour powered by the same indirect flight muscles. From these data, we estimate mass-specific energy expenditures and power outputs, and test how metabolic rates (MR) scale with body size (mass and intertegular span, ITS). Intertegular span is widely used as a reliable proxy for bee body size across taxa (28), facilitating comparisons with previous studies in bee energetics and morphology.

We hypothesised that: 1) the metabolic cost of floral buzzing would be at least as high as that of take-off, due to rapid and repeated activation of the IFMs; and 2) MR for both behaviours would scale positively with body mass and ITS, in line with biomechanical theory and prior studies showing that larger individuals expend more energy during high-frequency muscle use (29,30).

In addition, we examined intra-specific variation in metabolic performance by testing whether individuals differed consistently in their CO_2_ production across behaviours. While most metabolic studies focus on inter-specific variation, there is evidence that MR is a repeatable trait within individuals (31) that correlates with behavioural output (32), though the drivers of intra-specific differences remain poorly understood (33,34), particularly in insects (35,36). Variation may arise from developmental history (35), physiological maturation, or genetic background (34), and in social bees, colony-level differences may reflect heritable or epigenetic effects. To explore these possibilities, we estimated the repeatability (intra-class correlation coefficient, ICC) of CO_2_ production within individuals and tested whether colony identity explained variation. Understanding such variation is essential for evaluating the evolutionary constraints and flexibility of buzz pollination, an energetically demanding behaviour performed by some, but not all, bees.

## Materials and Methods

### Bees

We used three colonies of the buff-tailed bumblebee, *Bombus terrestris* (Biobest, Belgium), for experimental trials. Each colony had access to an *ad libitum* supply of Biogluc® ‘nectar’ solution provided within the colony box. Colonies were individually connected to flight arenas measuring 60 cm × 60 cm × 60 cm. These arenas were illuminated using overhead LED panels (595 mm × 595 mm, 3400 lumen; Aura Lunaria Dali DT8, Aura Light, Solna, Sweden) and maintained on a 9 h:15 h light:dark cycle. Ambient temperature in the laboratory ranged from 22–24°C, with relative humidity between 19–28%.

### Plants

Within each flight arena, bees were presented daily with eight inflorescences of *Solanum sisymbriifolium* Lam. (Solanaceae), a nectarless species that attracts and rewards pollinators solely through pollen. Like most species in the genus *Solanum*, *S. sisymbriifolium* possesses five poricidal anthers that release pollen when stimulated by bee-producing vibrations, i.e., it is a buzz-pollinated plant. *Solanum* is a well-established model for the study of buzz-pollination (19,21,37).

Seeds of *S. sisymbriifolium* were obtained from a commercial supplier (Chiltern Seeds, Wallingford, UK) and germinated following a published protocol (38). Plants were grown in the research glasshouses at Uppsala University under supplemental light 8h:16h dark:light, at 20°C during the day and 16°C at night.

For daily provisioning in the flight arenas, inflorescences were cut fresh each morning and placed in water-saturated Oasis IDEAL Floral Foam Maxlife (Oasis Floral Products, Houthalen-Helchteren, Belgium) held in plastic containers. For experimental trials, a single flower was selected and cut just below the calyx for insertion into the respirometry chamber.

### Experimental procedure

The respirometry chamber was a custom-built Perspex cylinder (10 cm length × 5 cm diameter), with a 3D-printed internal floor and sealed at each end using 3D-printed plugs secured by rubber bands (Fig. 1). Air scrubbed of CO_2_ and H_2_O was pumped through the chamber at a consistent rate of 400 ml/min regulated by mass flow controller (GFC17; Aalborg, NY, USA) positioned immediately prior to the test chamber. Air exiting the chamber was analysed for CO_2_ and H_2_O concentrations using an infrared LI-7000 CO_2_/H_2_O analyser (LI-COR Biosciences, Lincoln, Nebraska, USA), and for O_2_ concentration using an Oxzilla II dual-channel oxygen analyser (Sable Systems International, NV, USA). The chamber also contained an internal rotating door, 3D-printed to allow controlled bee access to the flower during trials (Fig. 1).

**Figure 1.**
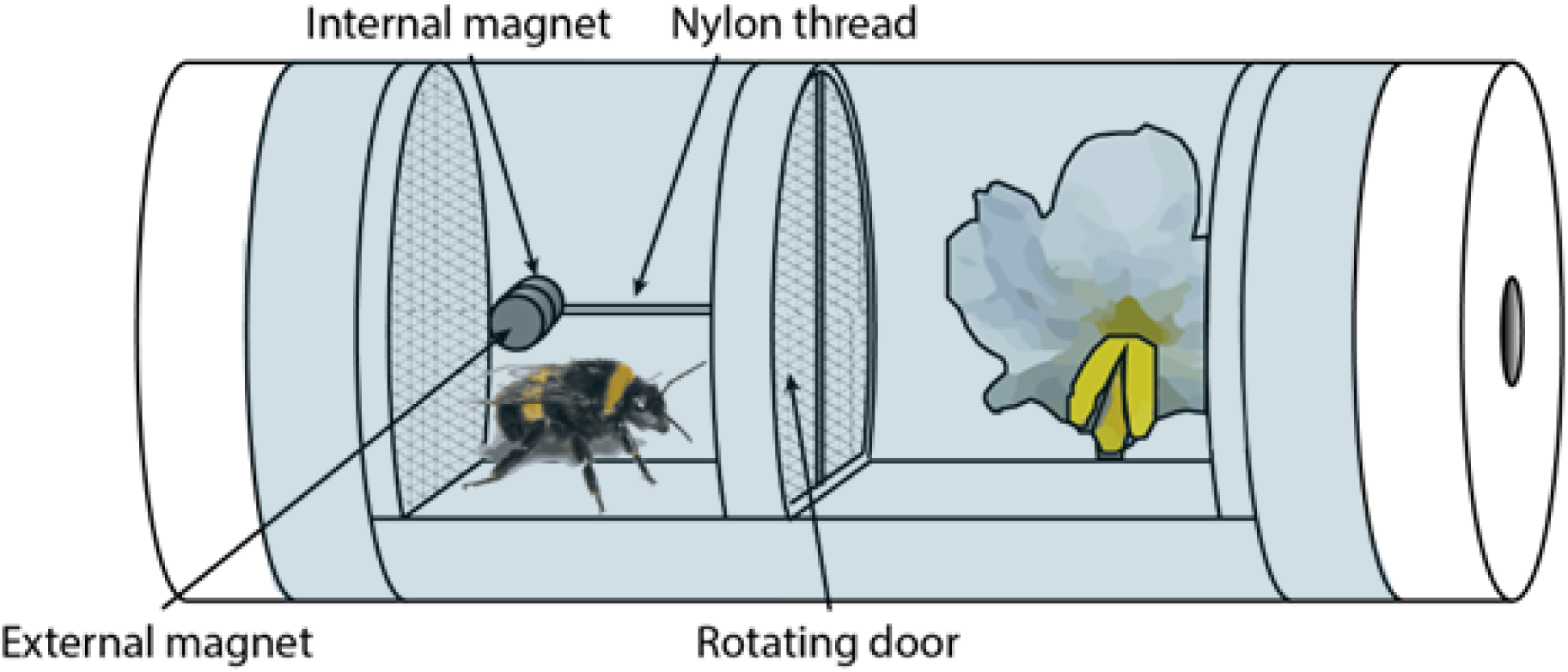
Schematic of the custom-built respirometry chamber used to assess bee gas exchange during floral interactions. The Perspex cylinder was divided into two compartments by a 3D-printed rotating door that allowed controlled access to a flower at the far end of the chamber. Bees were introduced into the front compartment and held for 3 minutes before the door was opened remotely using an external magnet. The door and magnet system, including a nylon-thread-mounted internal magnet, enabled non-invasive operation.

To initiate a trial, a group of five foraging bees were allowed access to the flight arena. When a bee was observed performing a floral buzz on a flower, it was captured using a queen catcher and weighed. A fresh flower was prepared by affixing a 2 x 2 mm piece of reflective tape to the back of one petal (to facilitate laser vibrometry recordings – see below for more details) and a small magnet (3 x 1 mm) to the base of the cut stem. A corresponding magnet inside the respirometry chamber ensured consistent flower placement across trials (Fig. 1).

Prior to the bee’s introduction to the chamber, baseline gas exchange was recorded with only the flower present until measurements stabilised. The bee was then introduced into the flower-free compartment of the chamber, and the internal door was kept closed for 3 minutes to allow gas clearance (Fig. 1). After this period, the door was opened remotely using an external magnet, allowing the bee to interact with the flower for 5 minutes. Once the bee returned to the flower-free compartment, the door was closed again, and post-interaction behaviour was recorded for an additional 5 minutes. The bee was then removed, re-weighed, euthanised by freezing at –20°C for at least 30 minutes, and photographed under a dissecting microscope. Image J was then used to measure inter-tegular span. Body mass was calculated as the mean of the pre– and post-trial weight.

### Recording of floral vibrations

We used a laser vibrometer to record floral vibrations (Portable Laser Vibrometer VGO-200, Polytec, Waldbronn, Germany). The laser provided contactless measurements of floral vibration velocity. The vibrometer was positioned approximately 20 cm in front of the flower and the laser trained on the retroreflective tape in the back using a mirror (broadband mirror 400-750 nm, Thorlabs, Molndal, Sweden).

The vibrational data were acquired and digitised with a C-Series Sound and Vibration module (9250; NI, Austin Texas, USA) and Compact DAQ chassis (CDAQ 9171; NI) connected to a laptop. Data were sampled at 10,240 samples per second with a 5 kHz low-pass filter and a maximum velocity range of 200 mm/s using a custom LabVIEW 2025 (NI) program. Data were saved as TDMS files to speed up data acquisition and exported with a second LabVIEW program as TXT files for downstream analysis.

Each trial lasted approximately 20 minutes. Vibrometry recordings were synchronised with respirometry and behavioural video data by manually obstructing the laser beam five times within the camera’s field of view prior to bee and flower introduction with respirometry recording initiated on the first obstruction.

### Data collection and analysis

In total, we collected data from 31 bees across three colonies. Ten of these were excluded from the analysis: eight because only one of the two behaviours of interest (buzzing and take-off) were extracted during clean event extraction (see below), and two because fewer than two events of one of the behaviours were available. This resulted in a final dataset comprising 21 bees (five, eight and eight per colony). An average of 12 floral buzzes and ten take-offs per bee were analysed, yielding a total of N = 260 floral buzzes and N = 213 take-offs.

#### Acoustic signal processing and buzz detection

The TXT files with vibration data were analysed in R version 4.4.3 (39) using the packages *seewave* (40) and *tuneR* (41) and code developed to extract and characterise bee buzzes (42). Vibration signals were high-pass filtered at 20 Hz using a Hanning window (length = 512) to reduce low-frequency background noise. Buzz events were automatically identified using the *timer* function from the *seewave* package (40), with a threshold set to 1.75% and a minimum duration of 0.1 s. For each detected buzz, acoustic features were extracted, including dominant spectral frequency, mean and median cepstral fundamental frequency, root mean square (RMS) amplitude, and both smoothed and raw peak amplitudes.

Following detection, a Python-based filtering pipeline was applied to ensure only biologically relevant floral buzzes were retained (43). First, buzzes were filtered based on their cepstral fundamental frequency; only events in the previously reported floral-buzz range (250-450 Hz) were retained (22). This removed non-floral signals such as artifacts, incidental vibrations or grooming behaviour. Second, buzzes were temporally filtered to retain only those that occurred during the floral interaction period, defined as the time between opening and closing the rotating door inside the respirometry chamber, which granted the bee access to the flower. Buzzes detected outside of this window were discarded. Because the laser vibrometer was aimed at a reflective strip attached to the back of the flower, only buzzes physically transmitted through the flower (i.e. true floral buzzes) could be detected. This ensured that only buzzes made in contact with the flower were included. Filtered buzzes were exported as individual uncompressed sound files (WAV). Each was manually verified through auditory inspection and labelled as either a true floral buzz or false positive. Only verified floral buzzes were retained for further analysis.

#### Behavioural annotation and synchronisation

To align behavioural, acoustic, and physiological data streams, a Python script (43) was used to detect the first peak in the respirometry trace corresponding to a series of five laser pokes. This peak marked the start of synchronisation and enabled alignment of the respirometry signal with acoustic and video recordings. Videos were manually annotated in BORIS (Behavioural Observation Research Interactive Software) (44) to identify take-off events and record the timestamp of the first laser obstruction. Take-off was defined as any wing-driven behaviour that lifted the bee’s body off all supports, ending when all legs contacted a surface again; due to the small chamber size, bees could not engage in sustained free flight. This operational definition is consistent with kinematic criteria used to demarcate take-off in insect studies (e.g. onset of centre-of-mass rise and loss of leg contact) (45).

#### Metabolic rate calculation

Gas exchange data were processed using ExpeData software (Release 1.9.27; Sable Systems Europe, Berlin, Germany). Bee-associated measurements were corrected by subtracting flower-only baseline values. Clean behavioural events (buzzing or take-off) were identified in the CO_2_ respirometry trace using a custom Python script (43). Each CO_2_ peak was matched to observed behaviours, and only those segments that corresponded exclusively to a single behaviour (either buzzing or take-off, but not both) were retained for analysis.

CO_2_ production (V̇CO_2_, in mL/h) was calculated by integrating the CO_2_ signal (converted from % to ppm) over time using the trapezoidal rule, a numerical method that estimates the area under the curve by summing small trapezoid-shaped sections. A 1-second buffer was applied to each event window to compensate for the mismatch between the rapid behavioural events and the slower response time of the LI-COR gas analyser. The integrated CO_2_ signal was corrected for flow rate (400 mL/min) and event duration and scaled to an hourly rate. To account for variation in ambient temperature, V̇CO_2_ values were normalised using a Q_10_ correction (Q_10_ = 2), with 23.3 °C used as the reference temperature, the average temperature across all trials. Mass-specific metabolic rate (MSMR) (mL/g/h) was obtained by dividing V̇CO_2_ by individual bee body mass.

#### Respiratory Quotient calculation

To estimate the energetic cost of individual buzzing and take-off events, we integrated the CO_2_ signal over the duration of each behavioural event. Because the response time of the oxygen analyser (Oxzilla II) was too slow (∼5–7 s) to reliably capture O_2_ dynamics during sub-second behaviours, we inferred oxygen consumption during buzzing (V̇O_2_) indirectly using the respiratory quotient (RQ):

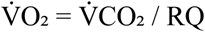

RQ was calculated for each bee from a continuous ∼5-minute recording corresponding to the floral interaction phase, defined as the time between opening and closing the rotating door of the respirometry chamber. During this interval, bees were allowed to freely interact with the flower. V̇O_2_ values for each event were then estimated by combining event-level CO_2_ MR (mL/h) with individual RQ values obtained from this floral interaction window.

#### Energy expenditure calculation

We estimated the energy equivalent of O_2_ consumption using a published conversion (46):

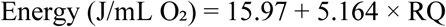

Total energy expended per event (in joules) was calculated as:

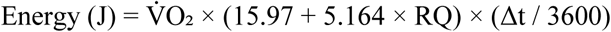

where Δt is the event duration in seconds. Mass-specific energy expenditure (J/kg) was calculated by dividing total energy by bee mass. Mass-specific metabolic power (W/kg) was then calculated by dividing total energy by bee mass and event duration:

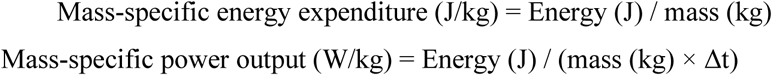

All MR were corrected to standard temperature and pressure, dry (STPD) conditions prior to these calculations, using procedures implemented in Python (43). This approach produced behaviour-specific energy estimates that accounted for individual physiological variation, body mass, and event duration.

#### Statistical analysis

All analyses were conducted in R (v4.4.3) (39). Unless stated otherwise, models included behaviour (buzzing vs. take-off) and colony identity as fixed effects, and bee identity as a random intercept to account for repeated measures. Model assumptions were assessed using the *DHARMa* package (47), and statistical significance was defined as *p* < 0.05 (two-tailed). Type II or III Wald tests (*car*) (48) (*lmerTest*) (49) were used to evaluate fixed effects.

We analysed absolute MR (V̇CO_2_, mL/h) and MSMR (mL CO_2_/g/h) at the event level, using linear mixed-effects models (*lmer*, *lme4*) (50). Response variables were log-transformed to meet model assumptions. Estimated marginal means (EMMs) were calculated using the *emmeans* package (51) to obtain back-transformed values, and pairwise comparisons were adjusted using the Kenward–Roger approximation. To assess the consistency of individual metabolic performance, we analysed event-level data using linear mixed-effects models with BeeID was included as a random intercept. This structure allowed us to partition variance into within– and between-individual components, and to calculate intraclass correlation coefficients (ICCs) using the *performance* package (52). ICCs quantify the proportion of total variance attributable to individual differences and were computed for the full dataset and separately for each behaviour (buzzing and take-off). For overall repeatability, values matched those obtained directly from the variance components of the full model. Behaviour-specific ICCs were obtained from models fitted separately for each behaviour. To determine whether colony-level variation in MSMR could be attributed to body size, we extended the model to include body mass and intertegular span (ITS) as covariates.

To investigate scaling relationships and behaviour-specific MR, we used individual-level summary data. For each bee, we calculated the mean log-transformed MR separately for buzzing and take-off. These per-individual means were modelled as a function of log-transformed body mass or intertegular span (ITS), behaviour, and colony identity using linear mixed-effects models (*lmer, lme4*) (50). Bee identity was included as a random intercept to account for repeated measures across behaviours. A similar model structure was used to assess scaling with ITS. Non-significant interactions between behaviour and the scaling variable were removed.

We compared mass-specific energy expenditure (J/kg) and mass-specific power output (W/kg) between behaviours using generalised linear mixed models (GLMMs) with a Gamma distribution and log link (*glmmTMB*) (53). EMMs and pairwise comparisons were derived using *emmeans* (51).

Differences in event duration between behaviour types was modelled using a Bayesian lognormal mixed-effects model implemented in the *brms* package (54), with behaviour and colony identity as fixed effects and bee identity as a random intercept. Posterior predictive checks confirmed that model fit was adequate.

To estimate the nectar volume (µL) required to fuel individual buzzing and take-off events, we fitted linear mixed-effects models (*lmer*) (50) with flower species, behaviour, body mass, and their two-way interactions as fixed effects. Colony identity was included as a fixed effect, and bee identity as a random intercept. Three flower species were included: *Rubus idaeus* (raspberry; 44.6% sugar, 8.37 J/µL) (55), *Helianthus annuus* (sunflower; 50.0%, 9.58 J/µL) (56), *Borago officinalis* (borage; 60.0%, 12.40 J/µL) (57). These species were selected because they are commonly visited by *Bombus terrestris* and have published nectar sugar concentration data. We converted sugar concentration to energy content using updated coefficients for floral nectar energetics (58). Nectar volume estimates were derived by dividing individual-level energy expenditure by species-specific energy content.

Post hoc comparisons were adjusted using Tukey’s method. Figures were created using *ggplot2* (59) and combined using *patchwork* (60). All codes are available on GitHub (43).

## Results

### Metabolic rates and respiratory quotients

Absolute MR (V̇CO_2_, mL/h) differed significantly according to behaviour (*F*_1, 457.03_ = 25.67, *p* < 0.001). During take-off, bees had a significantly higher MR than during buzzing (take-off EMM±SE: 11.2 ± 0.60 mL CO_2_/h, buzzing EMM±SE: 10.2 ± 0.55 mL CO_2_/h; *t* = –5.06, *p* < 0.001), equating to an approximate 9% increase in MR (*buzz / take-off* ratio = 0.915, 95% CI = [0.885, 0.945]). Colony identity did not significantly influence MR (*F*_2, 18.08_ = 1.33, *p* = 0.288).

MSMR was also significantly higher during take-off than buzzing (*F*_1, 461.39_ = 26.76, *p* < 0.001; take-off EMM±SE: 55.7 ± 2.23 mL CO_2_/g/h, buzzing EMM±SE: 50.9 ± 2.03 mL CO_2_/g/h; *t* = –5.17, *p* < 0.001), with a similar 8.6% increase in MSMR (*buzz / take-off* ratio = 0.914, 95% CI = [0.885, 0.944]). In contrast to absolute MR, MSMR varied significantly between colonies (*F*_2, 18.14_ = 13.34, *p* < 0.001). To assess whether this effect reflected underlying mass differences, we analysed body mass at the individual level. Body mass differed significantly between colonies (*F*_2, 18_ = 4.16, *p* = 0.033), with bees from Colony 4 being the lightest (mean ± SD: 0.18 ± 0.04 g, *n* = 8), Colony 3 intermediate (0.20 ± 0.02 g, *n* = 5), and Colony 5 the heaviest (0.23 ± 0.04 g, *n* = 8). A Tukey post hoc test confirmed that bees from Colony 5 were significantly heavier than those from Colony 4 (*p* = 0.028). To determine whether weight and size differences explained the colony effect on MSMR, we included both body mass and intertegular span (ITS) as covariates in the model. Colony identity remained a significant predictor (*F*_2, 16.04_ = 6.31, *p* = 0.010), while neither body mass (*F*_1, 15.45_ = 0.83, *p* = 0.376) nor ITS (*F*_1, 15.70_ = 0.23, *p* = 0.635) explained significant variation.

Individual bees were consistent in their MR across repeated behaviour events. The proportion of variance explained by differences between bees was high for absolute MR (ICC = 0.76 overall), with similar repeatability observed during buzzing (0.76) and even greater consistency during take-off (0.86). Repeatability for MSMR was somewhat lower (overall ICC = 0.63), but still substantial, with values of 0.66 for buzzing and 0.78 for take-off. These results indicate that individual differences in metabolic traits are repeatable, particularly during take-off.

The RQ was calculated for each bee over a five-minute window during which bees were allowed to interact freely with the flower. This period encompassed a range of behaviours, including buzzing and take-off. The mean RQ across all individuals was 0.940 ± 0.035 SEM (*n* = 21), with values ranging from 0.669 to 1.428 (range = 0.759). While most bees had RQ values close to 1.0, indicating primarily carbohydrate-based metabolism, a few individuals exhibited lower or higher values, suggesting mixed substrate use or transient non-steady-state metabolic conditions.

### Metabolic scaling

To assess the scaling of MR with body mass or size, we analysed the average MR of individuals for each behaviour (buzzing vs. take-off) (Fig. 2A). MR increased significantly with body mass (*F*_1, 17_ = 13.68, *p* = 0.002) and varied significantly between colonies (*F*_2, 17_ = 8.38, *p* = 0.003). The estimated scaling exponent for body mass was 0.82 ± 0.22, and the overall difference in MR between behaviours was statistically significant (*F*_1, 20_ = 4.48, *p* = 0.047), with take-off associated with higher values than buzzing (take-off EMM ± SE: 2.40 ± 0.045 mL CO_2_/h, buzzing: 2.32 ± 0.045 mL CO_2_/h; *t* = –2.12, *p* = 0.047).

**Figure 2.**
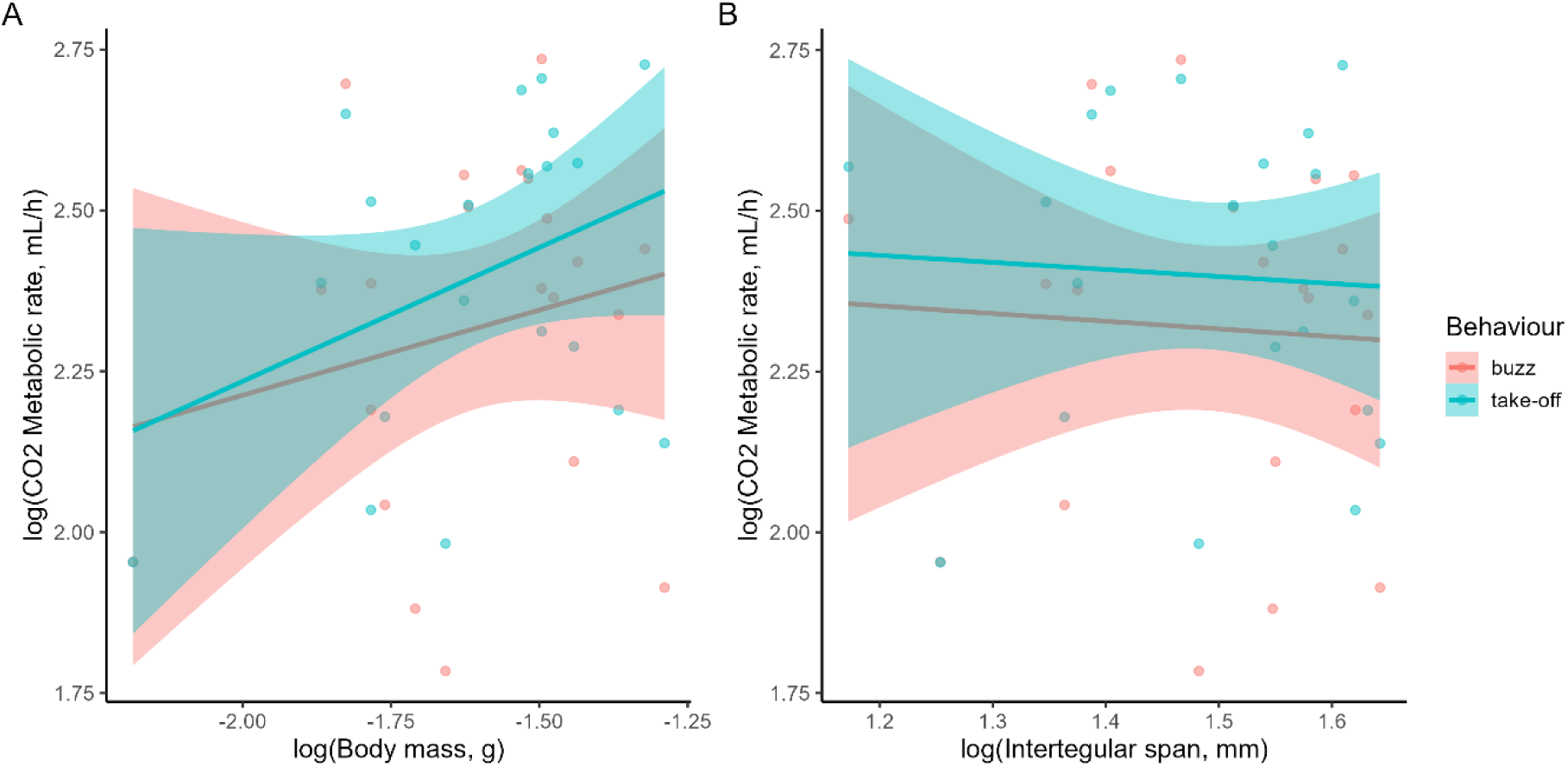
Metabolic scaling of CO_2_ production during buzzing and take-off behaviours. (A) Relationship between log-transformed CO_2_ MR and log-transformed body mass. (B) Relationship between log-transformed CO_2_ MR and log-transformed intertegular span (ITS). Coloured lines show fitted regression lines for buzzing (pink) and take-off (blue), with shaded areas representing 95% confidence intervals.

We also tested whether MR scaled with ITS, a proxy for structural body size (Fig. 2B). ITS did not significantly predict MR (*F*_1, 17_ = 2.92, *p* = 0.106). There was a significant effect of behavioural event (*F*_1, 20_ = 4.48, *p* = 0.047), and no statistically significant effect of colony identity (*F*_2, 17_ = 3.49, *p* = 0.054).

### Energetic and ecological costs

There was no effect of behaviour on mass-specific energy expenditure (Fig. 3A; χ²₁ = 1.20, *p* = 0.274), with bees expending a similar amount of energy during both buzzing and take-off (EMM = 0.11 ± 0.10 SE on the log scale). On the response scale, predicted energy expenditure was 471 J/kg for buzzing (95% CI: 403–550) and 422 J/kg for take-off (95% CI: 360–494), reinforcing the similarity between behaviours. In contrast, mass-specific power output, which accounts for the duration of the behaviour, was significantly higher for take-off compared to buzzing (Fig. 3B; χ²₁ = 4.36, *p* = 0.037; EMM= – 0.078 ± 0.037 SE on the log scale). Back-transformed estimates indicated that power output was 293 W/kg during buzzing (95% CI: 265–324) and 317 W/kg during take-off (95% CI: 286–350), highlighting the greater instantaneous demand of take-off events. Event duration was significantly shorter for take-off events compared to buzz events (estimate = −0.09, 95% CI = [−0.18, 0.00]). Colony identity had a significant effect on both energy expenditure and power output (energy expenditure: χ²_2_ = 24.43, *p* < 0.001; power output: χ²_2_ = 34.68, *p* < 0.001), but not event duration.

**Figure 3.**
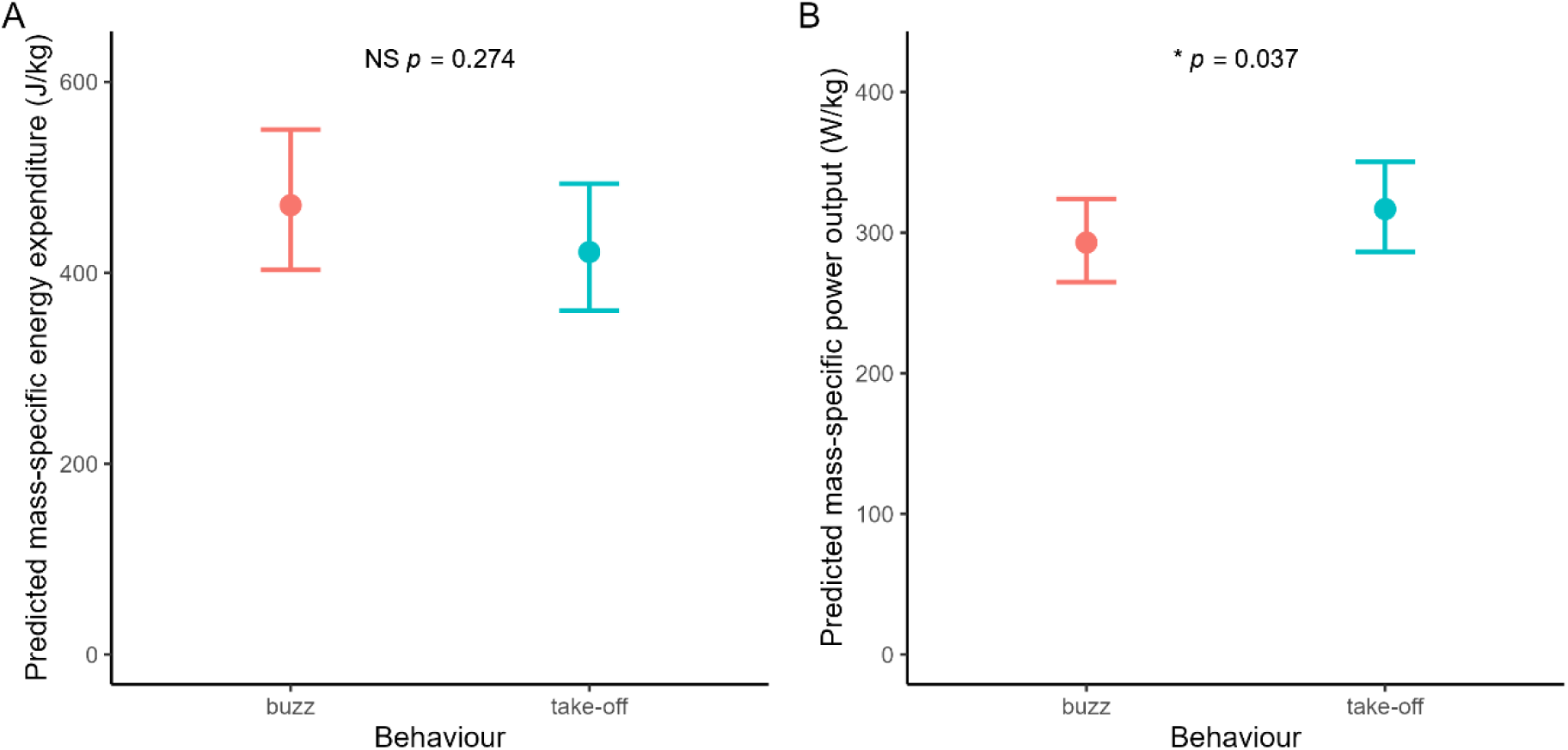
Predicted mass-specific energy expenditure and power output across behavioural contexts. (A) Estimated mass-specific energy expenditure (J/kg) and (B) mass-specific power output (W/kg) during buzzing and take-off behaviours. Values represent estimated marginal means ± 95% confidence intervals from generalised linear mixed models (GLMMs) with a Gamma distribution and log link, including colony identity as a fixed effect and individual identity as a random effect.

To evaluate the ecological cost of these behaviours, we estimated the nectar volumes required to fuel a single buzz or take-off event based on floral nectar compositions from three commonly visited plant species (Table 1). Flower species had a strong effect on nectar volume required (*χ²*_2_ = 36.47, *p* < 0.001), with borage requiring the least nectar and raspberry the most (borage < sunflower = raspberry; *p* < 0.001 for both comparisons). Behaviour also had a significant effect (*χ²*₁ = 4.50, *p* = 0.034), with buzzing requiring slightly more nectar than take-off (estimated difference = 0.0011 µL ± 0.0005 SE, *p* = 0.036).

**Table 1.**
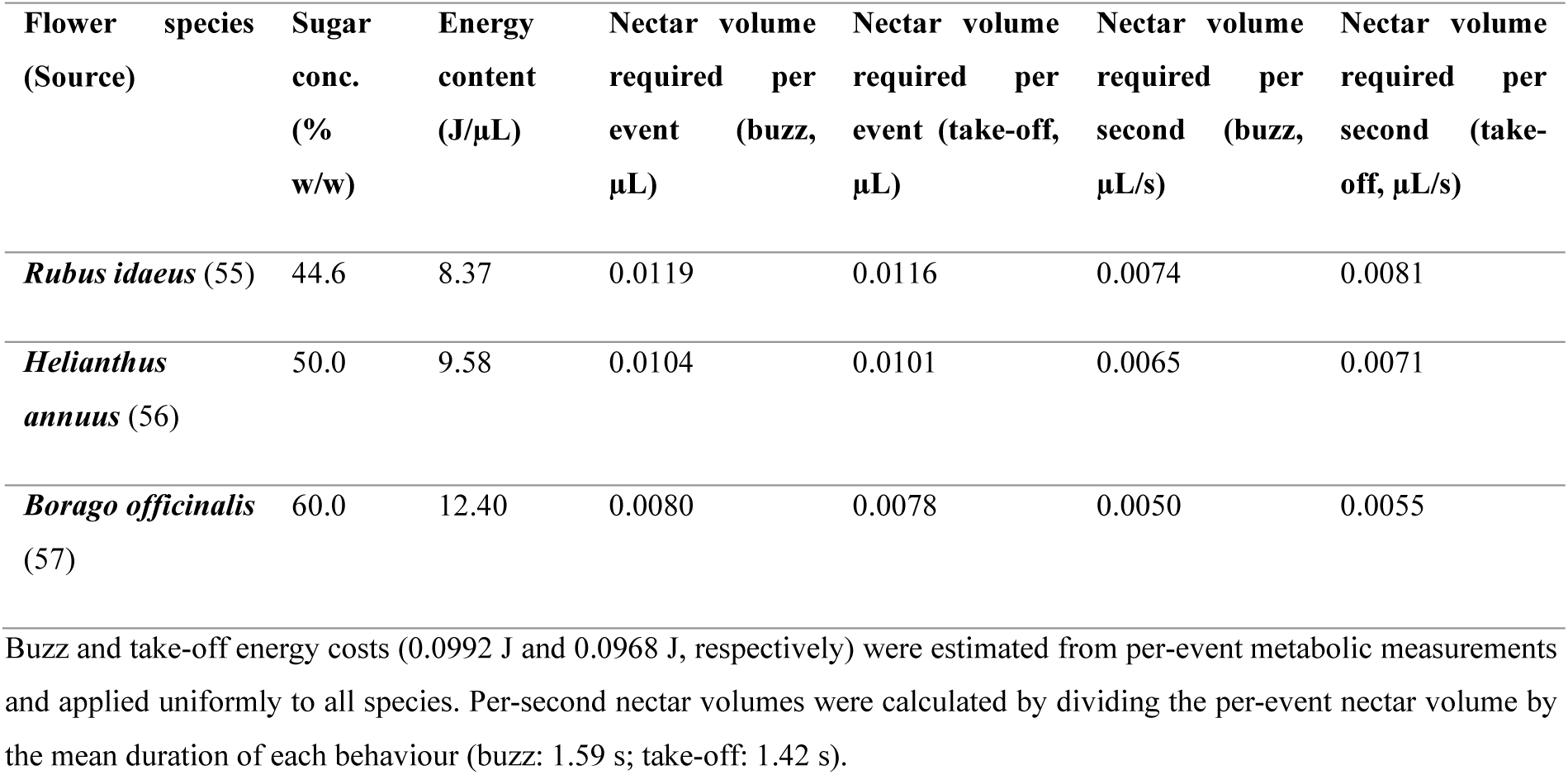
Estimated nectar volumes required to fuel a single buzz or take-off event, based on nectar composition from three flower species.

These effects reflect both variation in nectar sugar concentration among floral sources and the slightly longer duration of buzzing events. Colony identity also influenced nectar requirements (*χ²*_2_ = 10.80, *p* = 0.005), whereas body mass had no significant effect (*χ²*₁ = 0.74, *p* = 0.390). Overall, although energetic costs of individual buzz and take-off events were similar (∼0.10 J), the volume of nectar required to fuel them varied significantly with both behaviour and nectar quality.

Per-second nectar volumes (also shown in Table 1) were derived directly from the modelled power outputs and follow the same statistical patterns as those results; they are included for reference but were not analysed separately.

## Discussion

Our study provides the first direct measurement of the energetic costs incurred to bees during floral handling, specifically during buzz pollination. Because take-off is a well-characterised, high-power behaviour powered by the same indirect flight muscles, it offers a meaningful benchmark for interpreting the demands of buzzing. The comparison revealed that both behaviours are energetically costly, despite differing in their mechanical and temporal profiles. Take-off demanded more energy per unit time than buzzing; however, the longer duration of buzzing events resulted in similar total energy accumulated per behavioural event. This indicates that, over the course of a foraging bout, repeated buzzing could contribute as much to a bee’s total energetic expenditure as repeated take-offs. Such equivalence is important because it challenges the assumption that only flight imposes substantial energetic constraints, and suggests that floral buzzing may be a significant, previously under-appreciated component of foraging energy budgets. These results support our prediction that buzzing would impose a cumulative energetic cost comparable to that of take-off. We also predicted that MR would scale positively with body mass and intertegular span (ITS). While body mass was a significant predictor of MR across behaviours, ITS was not, offering only partial support for this hypothesis.

### Metabolic demands of take-off and buzzing: comparisons with prior studies

The high MR we observed during floral buzzing (10.2 mL CO_2_/h; 50.9 mL CO_2_/g/h) place it among the most energetically demanding behaviours recorded in *Bombus*. Resting workers, by contrast, typically show ∼0.3 mL CO_2_/h at 18 °C (61) and queens ∼7 mL CO_2_/g/h at 25 °C (62), underscoring a >30-fold elevation over resting metabolism in workers. Similar absolute rates have been reported for other intense behaviours: *B. impatiens* hovering or very slow forward flight at ∼9.5–11.7 mL CO_2_/h (63), and *B. lucorum* and *B. pascuorum* free flight at ∼50–70 mL O_2_/g/h (26), while our estimates exceed those for *B. lucorum* queens during flight (∼43 mL CO_2_/g/h) and brood incubation (∼27 mL CO_2_/g/h) (62). To enable direct comparison of our CO_2_-based rates with O_2_-based values, we measured RQ and found a mean of ∼0.94, consistent with carbohydrate-based metabolism and thus justifying CO_2_-O_2_ comparisons (64). While most RQ values were close to 1.0, deviations in either direction can occur in bees (65), and minor differences in analyser sensitivity could contribute to such deviations, although our 5-minute averaging window was intended to minimise this effect.

Compared to buzzing, take-off exhibited a ∼9% higher MR (11.2 vs. 10.2 mL CO_2_/h; 55.7 vs. 50.9 mL CO_2_/g/h). The observed differences in MR between behaviours likely reflect the distinct mechanical roles of the flight muscles during take-off and floral buzzing. In both behaviours, the same asynchronous indirect flight muscles (IFMs) are employed, but their functional output differs: during take-off, these muscles generate aerodynamic lift by actuating the wings, whereas in buzzing, the wings are decoupled, and muscle activity produces substrate-borne vibrations instead (14,66). Buzzing is therefore not a form of flight but rather a vibrational deployment of the flight apparatus. By eliminating the inertial and aerodynamic loads of the wings, bees can vibrate their thorax at higher frequencies, up to ∼300 Hz during floral buzzing in *B. terrestris*, compared to ∼150 Hz during flight (22). These higher frequencies have been hypothesised to carry a high energetic cost, especially when operating beyond the thorax’s natural resonance (66). Thus, buzzing has been considered to impose a unique form of physiological strain: high-frequency, non-resonant contractions that must be sustained over the entire duration of the behaviour. Our study found that while buzzing exhibited a lower MR than take-off on a per-second basis, the estimated mass-specific power outputs reached ∼317 W/kg during take-off and ∼293 W/kg during buzzing, confirming that both behaviours impose substantial muscular demands. These values are consistent with peak outputs recorded during forward flight or buzzing (26,67), versus ∼ 10 W/kg at rest (67). Importantly, despite take-off requiring higher instantaneous power, total energy expenditure per unit mass was statistically indistinguishable between behaviours, ∼471 J/kg for buzzing vs. ∼422 J/kg for take-off, reflecting the compensatory role of behaviour duration, and confirming the hypothesised cumulative energetic burden of sustained vibratory effort.

The consistency of our MR estimates with the upper range of reported values likely reflects both the instantaneous nature of our measurements and the specific intensity of the behaviours observed. Our measurements were taken at high temporal resolution and reflect peak activity, whereas some prior studies used averages over longer durations, potentially underestimating short bursts of intense effort. Similarly, differences in methodology, such as flow-through respirometry versus open-flow respirometry, calorimetry or kinematic modelling of power, can lead to variation in reported outputs, along with differences in species, castes, and sample sizes (68). Our power estimates align with the highest values found during forward flight or buzzing (26,67), suggesting that floral buzzing may push the flight musculature to its mechanical limits even in the absence of wing movement.

### Buzzing as a flexible behaviour

An interesting result from our repeatability analysis is the difference in the consistency of MR between the two focal behaviours. While take-off showed high intra-individual repeatability (ICC = 0.86), buzzing was less repeatable (ICC = 0.66), suggesting greater behavioural flexibility. This aligns with existing literature showing that bees might modulate buzz frequency, amplitude, and duration in response to floral traits, pollen yield, and prior experience (14,23). Buzzing performance is not constrained by biomechanical resonance in the same way as flight, allowing bees to adjust thoracic vibration characteristics beyond their natural wingbeat frequency (14,22). This flexibility likely underlies the greater metabolic variability we observed, as each buzzing event can be “tuned” to match the mechanical demands of a specific flower or foraging context. Take-off, in contrast, is a physically constrained, stereotyped action governed by the need to generate lift equal to body weight (69). Because biomechanical demands scale predictably with morphology, and because minimal lift cannot be compromised, take-off imposes a fixed energy threshold with limited scope for behavioural modulation. As such, metabolic performance during take-off is highly repeatable and tightly coupled to structural traits (69), consistent with our findings.

### Colony-level effects and metabolic variation

We observed significant colony-level differences in MSMR, power output, and energy expenditure, even after accounting for both body mass and structural size (measured as intertegular span, ITS). In contrast, absolute MR did not differ significantly between colonies, likely reflecting the dominant influence of body mass on total energy output. Body mass differed significantly across colonies, with workers from some colonies being consistently heavier than others. However, neither body mass nor ITS explained variation in MSMR, and colony identity remained a significant predictor even after controlling for both. This indicates that colony-level differences in metabolic performance are not driven solely by variation in body size or mass but likely reflect deeper physiological or developmental differences among colonies.

Previous work supports this interpretation: in *B. impatiens*, flight MR varied substantially among workers of similar mass owing to differences in wing morphology and muscle enzyme activity (70), traits that may also vary systematically across colonies. Although they did not detect a colony-level effect, other studies have reported consistent intercolony variation in traits such as worker lifespan (71) and behavioural flexibility (72), highlighting the biological significance of colony identity. In addition, colonies show heritable differences in morphological scaling relationships (73,74), suggesting that even subtle variation in developmental patterning could influence metabolic efficiency, independent of size. Taken together, these findings suggest that colony identity is a meaningful axis of physiological variation in bumblebees, likely shaped by genetic, maternal, or developmental factors. Future work should investigate how such factors influence muscle efficiency, thermoregulation, or metabolic regulation across colonies.

### Scaling patterns and predictors of metabolic rate

We observed significant scaling of absolute MR with body mass, with an estimated allometric exponent of 0.82 ± 0.22. This value aligns closely with previous estimates for flight MR in insects, particularly larger species (>10 mg), which scale around M^0.87^ (30) and reflects the elevated energetic demands of high-intensity behaviours such as take-off and buzzing. Behaviour also had a significant effect on MR, with take-off events associated with higher values than buzzing. However, the interaction between body mass and behaviour was not significant and was excluded from the final model, indicating a shared scaling relationship across behaviours.

Contrary to expectation, intertegular span (ITS), a common proxy for body size in bees (28), was not a significant predictor of MR. This suggests that morphological size alone does not account for the observed variation in energetic output. This may reflect the fact that ITS does not capture transient variation in traits such as thoracic muscle temperature, wingbeat frequency, or dynamic loading, which are more directly relevant to energetic performance during short, high-intensity behaviours (63,69,70). Assuming that body mass reflects not only structural size but also variation in nectar or pollen load (75), the observed scaling relationship may partly capture the increased energetic cost of carrying additional material. In this context, our results suggest that bees with greater body mass, due to loading, incur disproportionately higher metabolic costs during energetically demanding behaviours, even when engaging in the same activity. In such contexts, direct measurement of body mass offers greater predictive value.

### Ecological context and energetic trade-offs

Our results show that although the absolute energy expenditure of a single buzzing or take-off event was similar (∼0.10 J), buzzing required slightly more nectar to fuel the behaviour due to its longer duration. This arises because the same energetic cost is spread over a longer time, necessitating a greater volume of sugar solution to sustain muscular activity. Flower species significantly influenced the estimated nectar volume required, with borage needing the least and raspberry the most, reflecting differences in nectar sugar concentration.

These findings have important ecological implications. Buzz-pollinated flowers, such as those in the genera *Solanum* and *Cassia*, commonly offer abundant pollen but little or no nectar (14), requiring bees to expend muscular effort without an immediate source of fuel. Given the increased energetic demand we observed for buzzing, this presents a clear energetic challenge to foragers. Our data also imply that the grooming and pollen-packing phase following buzzing (76), adds further energetic cost due to its coupling with a second take-off. Together, these steps constitute a two-phase foraging sequence that increases both energetic expenditure and handling time.

We would therefore expect bees to adopt foraging strategies that mitigate these costs. Specifically, bees may supplement pollen collection during a foraging trip with nectar collection to maintain energy balance, particularly when visiting nectarless buzz-pollinated flowers. This prediction is supported by field observations (76), showing that bumblebees foraging on *Dodecatheon* alternated between buzzing for pollen and visiting nearby nectar-producing flowers. Similarly, honeybees carrying heavy loads adjusted their nectar intake depending on sugar concentration (77), suggesting regulation of fuel intake to task demand. In addition to modifying their foraging strategy, bees may also adjust the sequencing of behaviours within a foraging bout. Since the energetic cost of buzzing and take-off increases with body mass, it may be advantageous to perform energetically demanding behaviours early in a trip, such as buzzing for pollen, when overall load is low, and switch to nectar foraging later, when flight costs are higher. This kind of behavioural sequencing would allow bees to manage their energy budget more efficiently across a single event, consistent with models predicting that bumblebees optimise the ratio of pollen yield to energetic cost during foraging (78). More broadly, this strategy aligns with evidence that bees exhibit flexible foraging behaviour and adjust their decisions based on the relative cost and value of floral resources (13). Finally, bees may also rely on kinematic compensation to reduce energetic costs: when heavily loaded, bumblebees alter wing stroke amplitude and frequency to improve flight efficiency (63). Such adjustments could help offset the energetic penalties of load carriage during buzz pollination, though they may come with trade-offs in manoeuvrability or stability.

More broadly, our findings position buzz pollination as a highly specialised and energetically costly form of floral handling, likely to influence foraging decisions through effects on energy balance, trip structure, and floral choice (11,79). However, buzz-pollinated flowers represent only one end of a continuum of floral architectures and pollen presentation modes that may impose substantial handling costs on bees (80). These costs can vary with bee morphology, particularly body size and strength, which determine the ability to access rewards from mechanically complex flowers such as deep corollas, keel-type legumes, or joined anther cones (81–83). Handling efficiency may also be influenced by experience and learning, with complex floral morphologies often requiring longer handling times or repeated visits to achieve proficiency, thereby promoting flower constancy (9,10). Expanding the study of handling costs beyond buzz pollination to include a wider range of floral morphologies will be essential for understanding how these energetic and mechanical constraints shape foraging strategies, floral preferences, and plant–pollinator interactions across ecological contexts.

### Future directions and conclusions

This study opens several promising avenues for further research into the energetic ecology of bumblebee foraging. First, direct measurement of nectar and pollen loads during foraging bouts could clarify how dynamic changes in body mass influence energetic expenditure in real time, especially during transitions between buzzing, grooming, and flight. Integrating load quantification with metabolic measurements would offer mechanistic insight into how bees balance fuel intake with muscular effort across sequential behaviours.

Second, a deeper investigation into the biomechanical underpinnings of buzz pollination is warranted. Comparative analyses of wing loading, flight muscle architecture, and thoracic stiffness across individuals and colonies could reveal how bees differing in body mass, wing morphology, or muscle architecture achieve similar vibrational output and manage energetic efficiency. Examining muscle fibre type composition, mitochondrial density, and thoracic resonance properties may explain how some bees sustain high-frequency buzzing with lower energetic costs.

Third, the observed colony-level differences in metabolic traits suggest a strong developmental or genetic basis for variation in energetic performance. Future studies should investigate the physiological, genetic, and epigenetic factors underlying these intercolony differences. Experimental rearing under controlled conditions, coupled with transcriptomic or proteomic analyses, could help identify candidate genes or regulatory pathways influencing muscle efficiency, thermogenesis, and metabolic regulation.

Finally, field-based ecological studies are needed to contextualize these physiological findings within real-world foraging dynamics. Examining how bees sequence buzzing and nectar foraging, and how they modulate effort in response to floral diversity, reward distribution, and competition, would shed light on the energetic decision-making processes guiding pollinator behaviour. Observing foraging patterns in habitats dominated by nectarless, buzz-pollinated flowers could clarify the selective pressures shaping vibrational effort and floral preference.

In summary, our study provides the first direct comparison of metabolic costs associated with floral buzzing and take-off in bumblebees. While take-off imposes a higher instantaneous MR, the sustained duration of buzzing results in an energetically equivalent total cost per event. These findings confirm that floral buzzing imposes considerable muscular demands, despite the absence of aerodynamic lift, and highlight the significant energetic investment required for pollination of pollen-only flowers. Moreover, the variability in buzzing performance, both within and among colonies, suggests that bees adjust their vibrational effort in response to contextual cues such as floral traits, load status, or energetic reserves. This behavioural flexibility may allow bees to fine-tune their foraging strategy to maximize pollen gain while managing energy expenditure. Conversely, take-off remains a stereotyped, biomechanically constrained behaviour with high metabolic repeatability. By demonstrating that buzzing and take-off impose comparable energetic costs, our findings underscore the substantial physiological investment bees make in buzz pollination. This reinforces the view that pollen-only flowers exert strong selective pressures on pollinator foraging strategy and physiology. Future work linking vibrational effort to floral rewards, resource economics, and colony-level fitness will be critical to fully understanding the ecological and evolutionary dynamics of buzz pollination within plant–pollinator networks.

## Acknowledgments

We are grateful to Göran Arnqvist and Johanna Liljestrand Rönn for providing access to the gas analyser and advice for respirometry measurements. We thank Johan Fransson and Börje Söderling for help constructing the flight boxes, Fernando González-Almansa Laredo for assistance with initial bee measurements, Maria Uscka-Perzanowska for plant growth support, and the Vallejo-Marin Lab for support with plant and bee maintenance and discussions.

